# ISMapper: Identifying insertion sequences in bacterial genomes from short read sequence data

**DOI:** 10.1101/016345

**Authors:** Jane Hawkey, Mohammad Hamidian, Ryan R. Wick, David J. Edwards, Helen Billman-Jacob, Ruth M. Hall, Kathryn E. Holt

## Abstract

**Background:** Insertion sequences (IS) are small transposable elements, commonly found in bacterial genomes. Identifying the location of IS in bacterial genomes can be useful for a variety of purposes including epidemiological tracking and predicting antibiotic resistance. However IS are commonly present in multiple copies in a single genome, which complicates genome assembly and the identification of IS insertion sites. Here we present ISMapper, a mapping-based tool for identification of the site and orientation of IS insertions in bacterial genomes, direct from paired-end short read data.

**Results:** ISMapper was validated using three types of short read data: (i) simulated reads from a variety of species, (ii) Illumina reads from 5 isolates for which finished genome sequences were available for comparison, and (iii) Illumina reads from 7 *Acinetobacter baumannii* isolates for which predicted IS locations were tested using PCR. A total of 20 genomes, including 13 species and 32 distinct IS, were used for validation. ISMapper correctly identified 96% of known IS insertions in the analysis of simulated reads, and 98% in real Illumina reads. Subsampling of real Illumina reads to lower depths indicated ISMapper was reliable for average genome-wide read depths >20x. All ISAba1 insertions identified by ISMapper in the *A. baumannii* genomes were confirmed by PCR. In each *A. baumannii* genome, ISMapper successfully identified an IS insertion upstream of the *ampC* beta-lactamase that could explain phenotypic resistance to third-generation cephalosporins. The utility of ISMapper was further demonstrated by profiling genome-wide IS6110 insertions in 138 publicly available *Mycobacterium tuberculosis* genomes, revealing lineage-speific inserction and multi inserction hotspot.

**Conclusions:** ISMapper provides a rapid and robust method for identifying IS insertion sites direct from short read data, with a high degree of accuracy demonstrated across a wide range of bacteria.

## Background

Insertions sequences (IS) are small transposable elements that encode the proteins required for their own transposition. The ISfinder database [1] currently contains over 500 distinct IS. During transposition some IS create direct repeats, or target site duplications, in the sequences into which they are integrating. The presence and length of these duplications vary widely between IS and are characteristic of individual IS [2]. Rates of transposition vary between IS and host species, but are frequently in the order of the rate of nucleotide substitutions, making IS activity one of the more dynamic evolutionary forces at play in many bacterial genomes. The movement of IS can also have functional consequences for bacterial genomes. IS have been implicated in large changes to genome structure, by expanding in copy number in microbial genomes, with subsequent loss of IS resulting in inactivation of genes, pseudogene formation, mediating deletion of intervening sequences between two copies of the IS, or rearrangements of the genome [3].

In addition, IS insertions upstream of protein coding sequences can result in their enhanced expression, leading to different phenotypes depending on the function of the over-expressed gene. There are several known examples of IS-mediated gene expression leading to clinically important increases in antimicrobial resistance. For example, increased resistance to fluoroquinolones such as ciprofloxacin can result from the insertion of IS*1* or IS*10* upstream of the *acrEF* efflux pump in *Salmonella* Typhimurium [4], or the insertion of IS*186* upstream of the *acrAB* efflux pump in *Escherichia coli* [5]. In *Acinetobacter baumannii*, insertion of ISAba1 or ISAba125 upstream of the intrinsic beta-lactamase *ampC* can cause resistance to third generation cephalosporins including ceftazidime and cefotaxime [6, 7]. Insertions of the same IS in nearby locations can generate a composite transposon, capable of mobilizing the intervening sequence and transferring it to new genomic locations. For example, the composite transposon Tn*6168* was generated spontaneously via insertions of ISAba1 on either side of *ampC*, including one copy of ISAba1 that upregulates *ampC* expression [8]. Tn*6168* has then transferred into different *A. baumannii* backgrounds, conferring horizontally-acquired resistance to third generation cephalosporins [8].

IS insertions can also result in the upregulation of virulence genes in clinically important human pathogens. For example, an outbreak of tuberculosis in Spain in the 1990s was associated with the B strain of *Mycobacterium bovis* carrying an insertion of IS*6110* in the promoter region of the virulence gene *phoP*, resulting in its upregulation [9]. In *Neisseria meningitidis*, insertion of IS*1301* in the middle of the capsule locus has been shown to cause increased expression of operons on either side of the IS, contributing to protection from the human immune system and enhanced pathogenicity [10]. IS have also been shown to enhance niche adaptation in bacteria, for example IS*1247* insertion upstream of *dhlB* in *Xanthobacter autotrophicus* results in increased resistance to bromoacetate [11]. This region has also been mobilised by the IS and transferred to a plasmid [11]. In *E. coli*, IS*3* has been shown to up-regulate threonine expression, allowing the bacteria to adapt to a low-carbon environment and utilise threonine as its sole carbon source [12].

The profiling of IS insertion patterns has been used for typing purposes in numerous bacterial species of importance to human health. For example, copy number and position of IS*200* in *Salmonella enterica* [13], IS*6110* in *Mycobacterium tuberculosis* [14], IS*1004* in *Vibrio cholerae* [15] and ISAba1 in *A. baumannii* [16] has been used to profile these bacterial pathogens, allowing the identification and tracking of distinct subtypes. To date, IS-based typing schemes for various bacteria have relied on digesting the genome followed by either hybridizing IS probes to fragments in a gel or PCR probing [13–15]. The detection of precise insertion sites can be achieved using PCR, and may be done for typing purposes [17] or for the detection of functionally important insertions [7, 9].

With the advent of cheap high-throughput short-read sequencing, whole genome sequencing (WGS) of bacteria is increasingly common and is replacing traditional methods for characterizing and typing bacterial genomes. Unfortunately the detection of IS is complicated wherever read lengths are shorter than the length of the IS, as is the case for platforms that are currently most widely used – Illumina and Ion Torrent. IS insertion sites can readily be identified in finished bacterial genomes or in draft assemblies of genomes with single-copy IS, using tools such as nucleotide BLAST or ISfinder [1]. However where multiple copies of the same IS are present within a single genome (including on the chromosome and/or plasmids), this complicates assembly of short-read data and makes IS insertion sites difficult to identify reliably. The IS detection problem can be resolved using long-read sequencing technologies such as the SMRT Cell (Pacific Biosciences) or MinION (Oxford Nanopore) platforms; however given the relative cost efficiency and reliability of short-read sequencing, together with the current widespread use of Illumina for bacterial WGS and wealth of available short-read data for clinically important bacteria, there remains a need for a simple tool to identify IS insertion sites from short-read data.

Several studies report the use of mapping-based approaches to identify IS insertion sites from bacterial short-read data [18, 19], however none provide software code or validation of the approach used. There are tools available for detecting transposons or structural variation in genomes, for example MindTheGap [20] and BreakDancer [21], however these do not perform well in the identification of IS in bacterial genomes nor were they designed to do so. Some programs could potentially be used for this purpose, such as RelocaTE [22] and RetroSeq [23], however these require additional input or prior knowledge about the IS which may not always be available. TIF (Transposon Insertion Finder) [24] and *breseq* [25] could potentially be used for the detection of IS insertion sites in bacterial genomes, however they were not designed specifically for this purpose and did not perform well on our data sets (see Results).

Here we present a rapid and robust tool for accurate detection of IS sequences, including insertion site and orientation, direct from short-read data. The method is freely available in the form of open-source code called ISMapper, and here we validate its use via analysis of simulated and real short-read data from a range of IS and bacterial species. ISMapper requires short reads and query IS sequences as input, and can be used either for typing against a reference genome or to assist with manual resolution of complex short-read assemblies.

## Implementation

An overview of the ISMapper workflow is shown in Figure 1. ISMapper takes as input: (i) a set of paired end Illumina reads for an isolate of interest, (ii) an IS query sequence in fasta format, and (iii) either a reference genome (for typing) or an assembly of the read set (for assembly improvement), in GenBank or FASTA format (Figure 1a). Paired end Illumina reads are mapped to the IS query sequence using BWA-MEM (v0.7.5a or later) [26]. From the resulting alignment file (SAM format), unmapped reads whose pairs map to the end of the IS query sequence (that is, reads representing the sequences directly flanking the IS) are extracted using SAMtools view (v0.1.19 or later) [27] to retrieve reads based on SAM flags (Figure 1b). Specifically, left flanking reads (taking input sequence as left to right) are extracted using flag ‘–f 36’ and right flanking reads are extracted using flag ‘–F 40 –f 4’ and stored in separate BAM files, which are then converted to FASTQ format using BedTools (v2.20.1) [28]. In addition, Samblaster (v0.1.21) [29] is used to extract from the SAM file any reads that map to the end of the IS and extend into the neighbouring sequence (i.e. “soft clipped” reads, Figure 1b). The resulting FASTQ file is filtered using BioPython to extract the soft clipped portion of reads, where those sequences fit a specified size range (default 5-30 bp). The resulting sequences are sorted into left and right flanking sequences; these are each mapped separately to the reference genome or assembly using BWA-MEM, to identify the location(s) of the query IS in the genome under analysis (Figure 1c). Insertion site information is extracted from the resulting alignments using BedTools (coverage command) to summarise coverage of the reference by left and right flanking reads; these are filtered by read depth (default, minimum read depth ≥6x) to minimize false positive hits, and regions that overlap or are separated by a short distance (default, ≤100 bp) are merged using BedTools (merge command). Pairs of left and right flanking regions that likely represent either side of the same IS insertion are identified on the basis of positional information, using BedTools (intersect and closest commands). Left and right regions that overlap are considered to indicate a novel IS insertion not present in the reference, with the overlap resulting from target site duplication arising during IS transposition (Figure 1c, novel site). Where left and right regions are separated by a sequence that is approximately the length of the IS query, the intervening sequence is extracted and compared to the IS query using nucleotide BLAST+ (v2.2.25 or later) [30] to confirm whether this is a known insertion site that is present in the reference (Figure 1c, known site).

**Figure 1:**
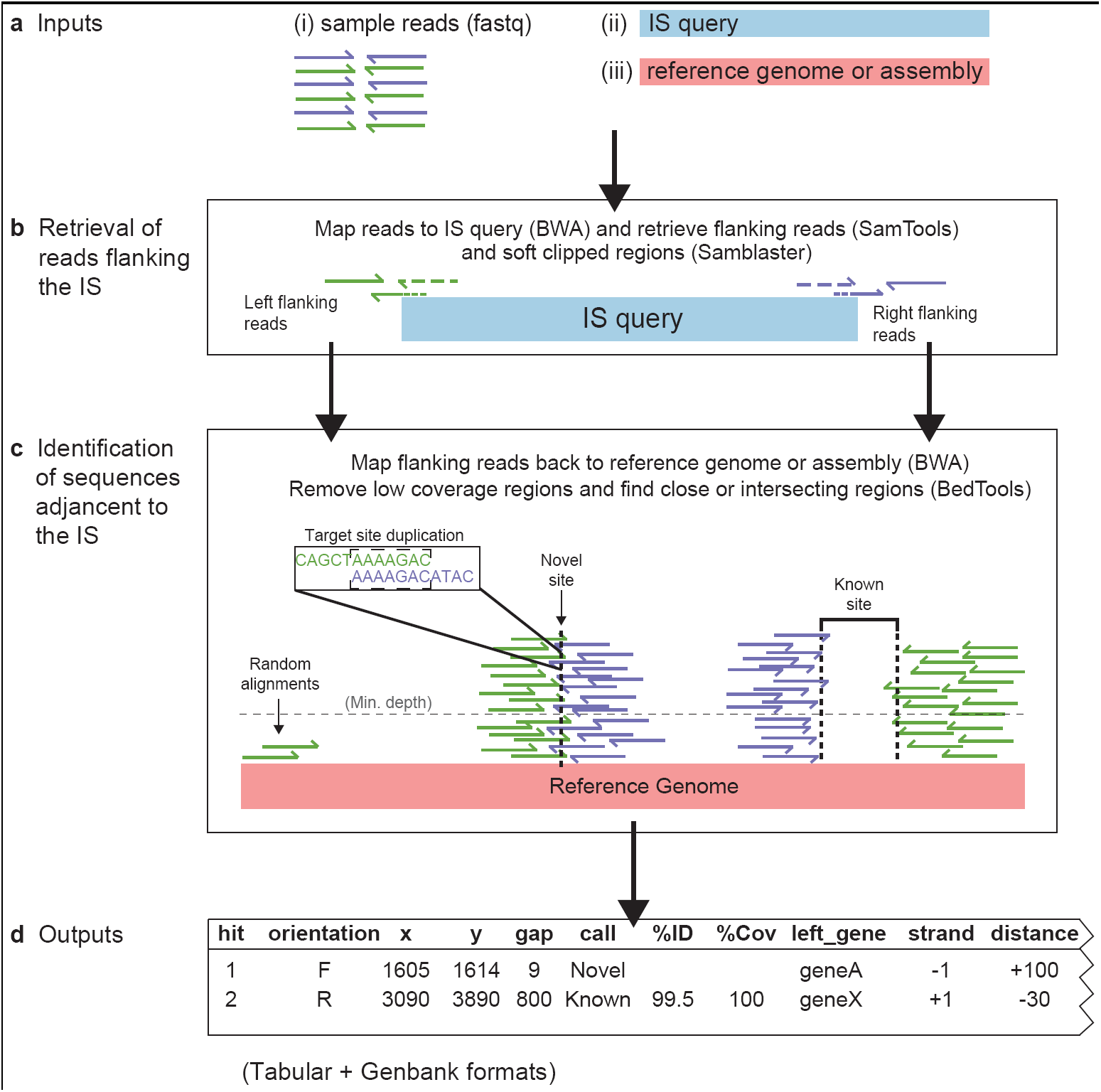
Workflow for ISMapper. (**a**) Inputs are reads (fastq format) and an IS query (fasta), as well as either a reference genome to compare to or an assembly of the reads (fasta or genbank). (**b**) Reads are mapped to the IS query using BWA (dashed lines) and their pairs (i.e. flanking reads, solid lines) are retreived from the resulting SAM file using mapping flags (SAMTools). The unmapped component of soft clipped reads (solid+dashed lines), identified from the SAM file using Samblaster) are also retrieved using BioPython. (**c**) Flanking and softclipped read sequences are then mapped to either the reference genome or the read assembly (BWA), and the final mapping output is then filtered for depth to remove low coverage regions. Left and right end blocks are extracted from the resulting BAM file (using BedTools) and compared to one another to find either intersecting regions, indicating a novel IS position, or close regions, indicating a known IS position in the reference. In a novel position, overlapping sequences from the left and right ends most likely indicate the target site duplication generated during IS transposition (zoomed in section). (**d)** Results are tabulated, indicating the position and orientation of each site, whether it is novel or known, and information about the genes flanking the insertion site.

Extensive testing of ISMapper revealed that it was sometimes unable to resolve IS positions that were adjacent to a repeat region (segments of DNA that were repeated multiple times around the genome; see Results). This is because when the IS-flanking reads were mapped back to the reference genome, those that belonged to the neighbouring multi-copy sequence were randomly assigned by BWA-MEM to the various locations of the repeat sequence, resulting in low read depth at the ‘true’ IS-adjacent copy of the multi-copy sequence, which can fall below the minimum depth filter (Figure 2). In such cases, the sequence on the other side of the IS is usually not a multi-copy sequence and thus does not suffer the same problem, and so is usually identified as a confident IS-flanking region without a corresponding partner region (Figure 2, purple block). Therefore, when ISMapper identifies an un-partnered IS-flanking region, it checks the original alignments for evidence of a nearby low-coverage partner region that failed to pass the depth filter and returns this as a potential but uncertain IS location, indicated by a ‘?’ character in the results table (Figure 2, green block).

**Figure 2:**
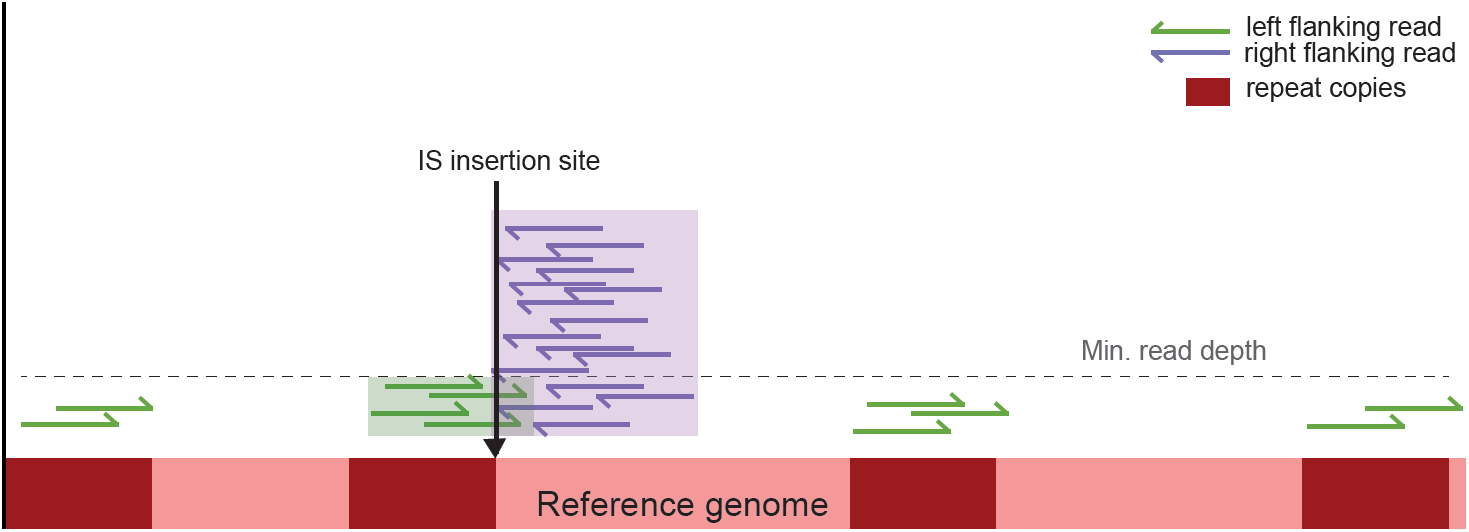
Issues can arise in the second mapping step of ISMapper if the IS insertion is next to a region that occurs more than once in the reference genome. In this example, the left flanking reads (green arrows) have mapped to all possible copies of the repeat sequence (dark red boxes) and are each randomly assigned to a repeat copy by BWA, resulting in each copy falling below the read depth cut-off (default 6x). To overcome this, where an unpaired flanking region is identified (purple box), ISMapper searches for a potential low-depth partner flanking region that is close to or intersecting the confident region (green box). This is recorded as an uncertain call, labelled in the output with the ‘?’ character.

ISMapper generates two main output files summarizing the results: (i) a GenBank file of the reference sequence, annotated with the IS-flanking regions and (ii) a table indicating the locations and characteristics of each IS-flanking region identified (Figure 1d). The table includes details of the location of the IS insertions; the distance between the left and right flanking regions (where a negative number indicates an overlap of left and right regions, indicating the size and sequence of the target site duplication); a call as to whether the insertion is present in the reference or is a novel insertion site (and, where the insertion site is present in the reference, the percent coverage and sequence homology with the IS query); and details of the gene(s) closest to the IS insertion site (including locus tag, product, gene name and distance from the IS to the start codon). Insertions are also marked to indicate less confident calls. A ‘*’ indicates an imprecise hit; i.e., where the gap between left and right regions is larger than expected for a novel insertion, but is not consistent with an IS insertion at that location in the reference. A ‘?’ indicates an uncertain hit, where only one end (left or right of the predicted insertion) passes the minimum read depth threshold; this often occurs when the IS is inserted within or adjacent to a multi-copy sequence, as described above (Figure 2). When run in assembly improvement mode, the table produced is simpler and indicates which contigs are predicted to end adjacent to the IS (indicating left or right orientation), assisting the user to decide whether some contigs could be joined together based on the available IS evidence.

ISMapper is lightweight code – a test run on a laptop computer (MacBook Air) with 8GB of RAM and a 1.3GHz i5 processor was able to analyse a read set comprising 2.5 million 100 bp paired-end reads in approximately ten minutes for a single IS query. Because ISMapper analyses each read set and query IS independently, screening of multiple read sets and query IS can be easily performed in parallel across multiple cores. To facilitate easy compilation of results from multiple jobs, ISMapper includes a Python script to cross-tabulate results from multiple read sets, generating a single summary table per query IS (script ‘compiled_table.py’).

## Results and Discussion

### Validation of IS detection using simulated reads

Nine publicly available genomes from a variety of bacterial genera, and including both chromosomes and plasmids, were downloaded from NCBI (Table 1). ISfinder [1] was used to identify the IS present in each genome sequence. All sequences that had >50% identity to a sequence in ISfinder and were present in at least two copies were extracted as query IS for testing with ISMapper. Nucleotide BLAST+ was used to confirm the precise locations and orientations for each query IS in all genomes (total 251 insertions of 17 distinct IS, see Table 1). Short reads (100 bp) were simulated from each genome sequence using the wgsim command in SAMtools (v0.1.19), with default parameter settings.

**Table 1:** Validation of ISMapper using simulated reads. * indicates ISMapper was unable to resolve some IS positions due to repeat regions.

| Isolate | Accession | IS | Found | Orientation |
| --- | --- | --- | --- | --- |
| *S. Typhi* CT18 | NC_003198 | IS200 | 26/26 | 26/26 |
|  |  | IS1 | 3/3 | 3/3 |
| *S. Typhimurium* LT2 | NC_003197 | IS200 | 6/6 | 6/6 |
|  |  | ISSty2 | 2/2 | 2/2 |
|  |  | IS1351 | 2/2 | 2/2 |
| *S. Typhi*<br>plasmid pHCM1 | NC_003384 | IS26 | 4/4 | 4/4 |
|  |  | ISVsa5 | 2/2 | 2/2 |
|  |  | IS1 | 5/5 | 5/5 |
| *S. Paratyphi*<br>plasmid pAKU_1 | AM412236 | IS26 | 6/6 | 6/6 |
|  |  | ISVsa5 | 2/2 | 2/2 |
|  |  | IS1 | 7/7 | 7/7 |
| *K. pneumoniae*<br>plasmid pNDMAR | JN420336 | IS3000 | 3/3 | 3/3 |
|  |  | IS26* | 3/5 | 3/5 |
|  |  | ISEcp1 | 2/2 | 2/2 |
| *Yersinia pestis* C092 | NC_003143 | IS100* | 43/44 | 43/44 |
|  |  | IS1661* | 7/8 | 7/8 |
|  |  | IS1541* | 61/64 | 61/64 |
| *Escherichia coli*<br>O104:H4 | NC_018658 | IS1 | 10/10 | 10/10 |
|  |  | IS421 | 4/4 | 4/4 |
|  |  | IS609 | 4/4 | 4/4 |
|  |  | ISEc1 | 4/4 | 4/4 |
|  |  | ISKpn26 | 4/4 | 4/4 |
| *E. coli* O157:H7 | NC_002695 | IS629* | 17/18 | 17/18 |
|  |  | IS609 | 2/2 | 2/2 |
|  |  | ISEc1 | 4/4 | 4/4 |
|  |  | ISEc8* | 9/10 | 9/10 |

ISMapper was run with default parameter settings on each combination of genome, query IS and simulated reads. ISMapper was able to accurately locate each IS position and its orientation (ranging between 2 and 61 positions per genome) for the majority of genomes (Table 1). In total, 96.4% of IS insertions were correctly detected. The exceptions occurred in three genomes (*K. pneumoniae* plasmid pNDMAR, *Y. pestis* CO92 and *E. coli* O157:H7), in which ISMapper correctly identified 151 IS insertion sites and failed to identify nine (94% detection). Closer inspection revealed that the missed IS were each located next to multi-copy repeat sequences, complicating the second mapping step as discussed above and outlined in Figure 2. Switching on reporting of all alignments above a mapping score threshold of 30 (-a and -T 30 in BWA-MEM) enabled the detection of a further IS*100* site in *Y. pestis*. By default this option is turned off in ISMapper as it tends to create noise in the mapping, making it more difficult to distinguish true and false positives; however this can be useful if an IS site of interest is known or suspected to be flanked by further repeats.

### Validation of IS detection using real Illumina read sets derived from isolates with finished genomes

Next we validated ISMapper using six genomes for which both Illumina read data and finished genomes were publicly available (Table 2). Each finished genome sequence was analysed with ISfinder [1] to identify query IS for testing as described above, and nucleotide BLAST was used to confirm the precise locations and orientations of each IS in each genome. The resulting test set comprised 106 insertions of 14 query IS. Using default settings, ISMapper was able to accurately identify each IS insertion site and its orientation, between 2 and 26 per genome, for the majority of genomes (Table 2). In total, 104 (98%) IS insertions were correctly detected by ISMapper. Three of four IS431mec insertions in Staphylococcus aureus TW20 were correctly detected, however the fourth was missed by ISMapper as it was flanked by another IS431mec and further repeat sequences. Two of three IS1 insertions in Salmonella Typhi CT18 were correctly detected however a third, located between tviE and tviD, was problematic. ISMapper was able to identify the region flanking the IS at tviE, but was unable to detect the corresponding region in tviD. Inspection of a BWA-derived alignment of the full Illumina read set to the CT18 chromosome showed that the entire region spanning from tviD to tviA was devoid of sequence reads, suggesting that this region may have been deleted during culture in the laboratory prior to the extraction of DNA for Illumina sequencing. This region encodes the biosynthesis of the Vi capsule of S. Typhi, and is known to be lost sporadically during culture [31]. This illustrates that situations where one end of the IS is detected but the other is not can often be ‘accurate’ in the sense that the result reflects underlying structural variation in the genome, including potentially IS-mediated deletions.

**Table 2:** Validation of ISMapper using real Illumina reads for which finished genomes were also available. * indicates ISMapper was unable to resolve some positions due to repeat regions.

| Isolate | Genome | FastQ | IS | Found | Orientation |
| --- | --- | --- | --- | --- | --- |
| *Streptococcus suis* P1/7 | AM946016 | ERR225612 | ISSu3 | 4/4 | 4/4 |
|  |  |  | ISSu4 | 2/2 | 2/2 |
| *Staphylococcus aureus* TW20 | NC_017331 | ERR043367 | ISSep3 | 3/3 | 3/3 |
|  |  |  | IS256 | 8/8 | 8/8 |
|  |  |  | IS431mec* | 3/4 | 3/4 |
|  |  |  | IS1181 | 2/2 | 2/2 |
| *Klebsiella pneumoniae* NJST258_1 | CP006923 | SRR1166975 | ISKpn1 | 6/6 | 6/6 |
|  |  |  | IS5B | 8/8 | 8/8 |
|  |  |  | IS903B | 2/2 | 2/2 |
|  |  |  | IS1294 | 3/3 | 3/3 |
|  |  |  | ISKpn18 | 2/2 | 2/2 |
|  |  |  | ISKpn26 | 7/7 | 7/7 |
| *S. Typhi* CT18 | NC_003198 | ERR343331 | IS200 | 26/26 | 26/26 |
|  |  |  | IS1* | 2/3 | 2/3 |
| *S. Typhi* Ty2 | AE014613 | ERR343332 | IS200 | 26/26 | 26/26 |

### Detection of antibiotic resistance-mediating IS insertions in *Acinetobacter baumannii*, confirmed by PCR

The genomes of seven ceftazidime resistant *A. baumannii* isolates, belonging to global clone (GC) 1, were sequenced via Illumina HiSeq to generate 100 bp paired end reads. Resistance gene screening of the Illumina data using SRST2 [32] and the ARG-Annot database [33] confirmed earlier PCR data indicating that none of these isolates carried acquired extended spectrum beta-lactamase (ESBL) genes that can confer resistance to third-generation cephalosporins. However, it is known that the insertion of ISAba1 upstream of the intrinsic chromosomally encoded *ampC* betalactamase gene can cause increased resistance to third-generation cephalosporins in *A.*baumannii* [6].*

We used ISMapper to screen for the ISAba1 query sequence (accession AY758396), sourced from ISfinder [1]. Using default parameters, ISMapper identified ISAba1 insertions in all seven GC1 genomes. IS positions were assessed relative to the genome sequence of *A. baumannii* GC1 reference A1 (accession CP010781). ISMapper found between 3 and 5 ISAba1 insertions in each GC1 isolate, including an insertion upstream of *ampC* in all 7 genomes that was in the orientation required to induce upregulation and explain the observed cephalosporin resistance phenotype (Figure 3). In addition, out of 29 total ISAba1 insertions, ISMapper was able to correctly identify 26 target site duplications (9 bp in the case of ISAba1). All ISAba1 insertions were novel compared to the reference genome A1 (Figure 3) and were confirmed using PCR, as described in [6].

**Figure 3:**
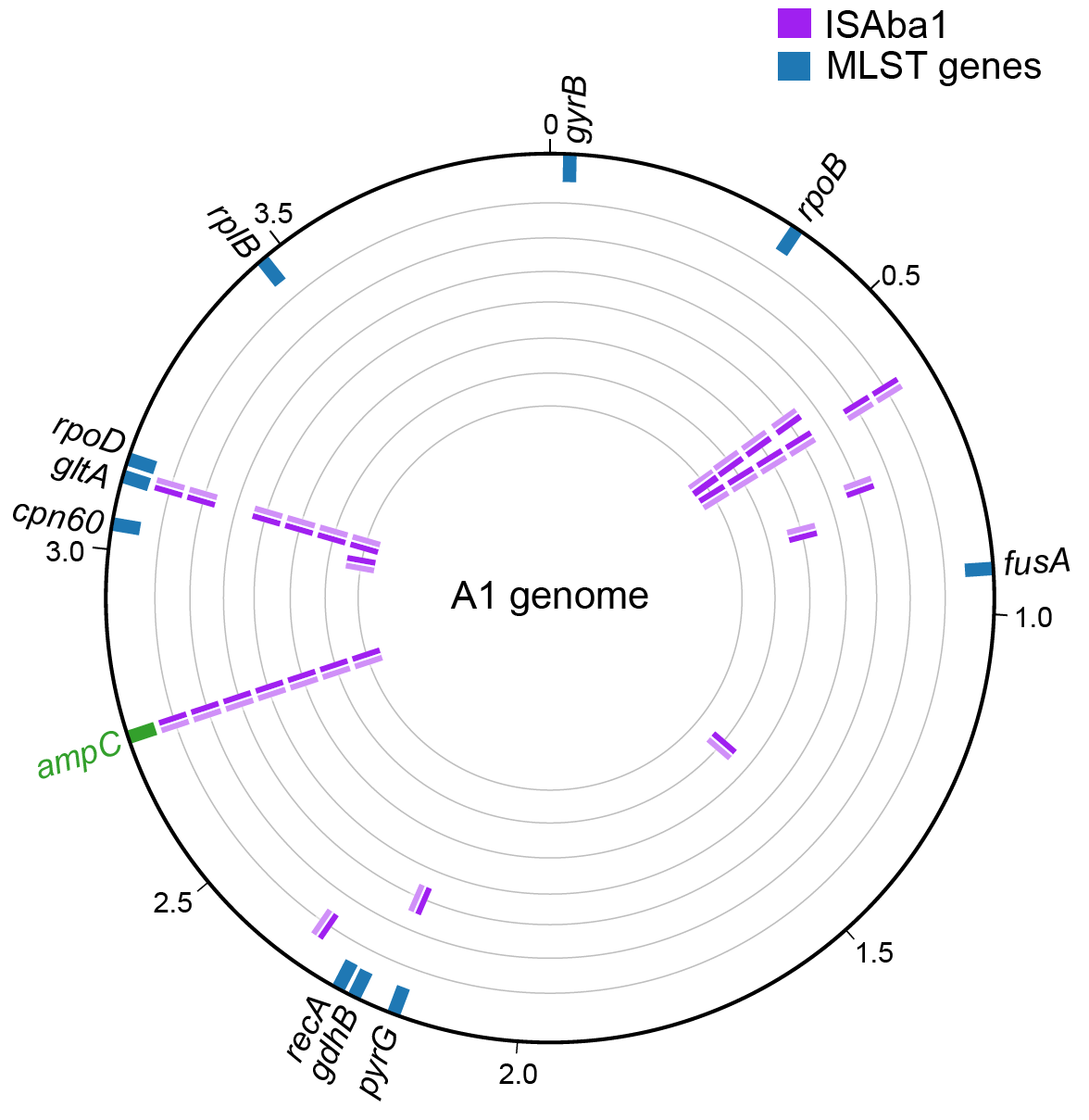
ISAba1 positions detected in seven *Acinetobacter baumannii* genomes. A1 reference genome is indicated by the outer black circle, tick marks indicate genome coordinates (in Mbp), genes targeted in the two available MLST schemes are marked in blue for orientation purposes. ISAba1 detected within the individual test genomes are shown on inner rings (purple), orientation of the IS is indicated by shading (dark, left end; light, right end). Novel IS insertions were identified upstream of the beta-lactamase *ampC* (green) in all isolates, explaining the observed phenotypic resistance to third generation cephalosporins.

### Impact of read depth on ISMapper performance

To test the effect of read depth on the performance of ISMapper, each of the seven GC1 *A. baumannii* read sets were randomly subsampled to depths of approximately 10x, 15x, 20x, 25x, 50x, 75x and 100x, with ten replicates per depth level per read set.ISMapper was then run using default settings to screen for ISAba1 insertions. The results indicated that at mean genome-wide read depths of approximately 20x, ISMapper was able to identify 95% of insertions correctly (Figure 4). However, all of these calls were either imprecise (gap size larger than expected) or uncertain (high coverage end paired with a low coverage end). An average genome-wide read depth of ~50x was required to find all insertions, with confident calls for >60%, however there was clearly some variation depending on read quality (Figure 4). To achieve 100% detection with high confidence, average genome-wide read depths of >75x were required (Figure 4).

**Figure 4:**
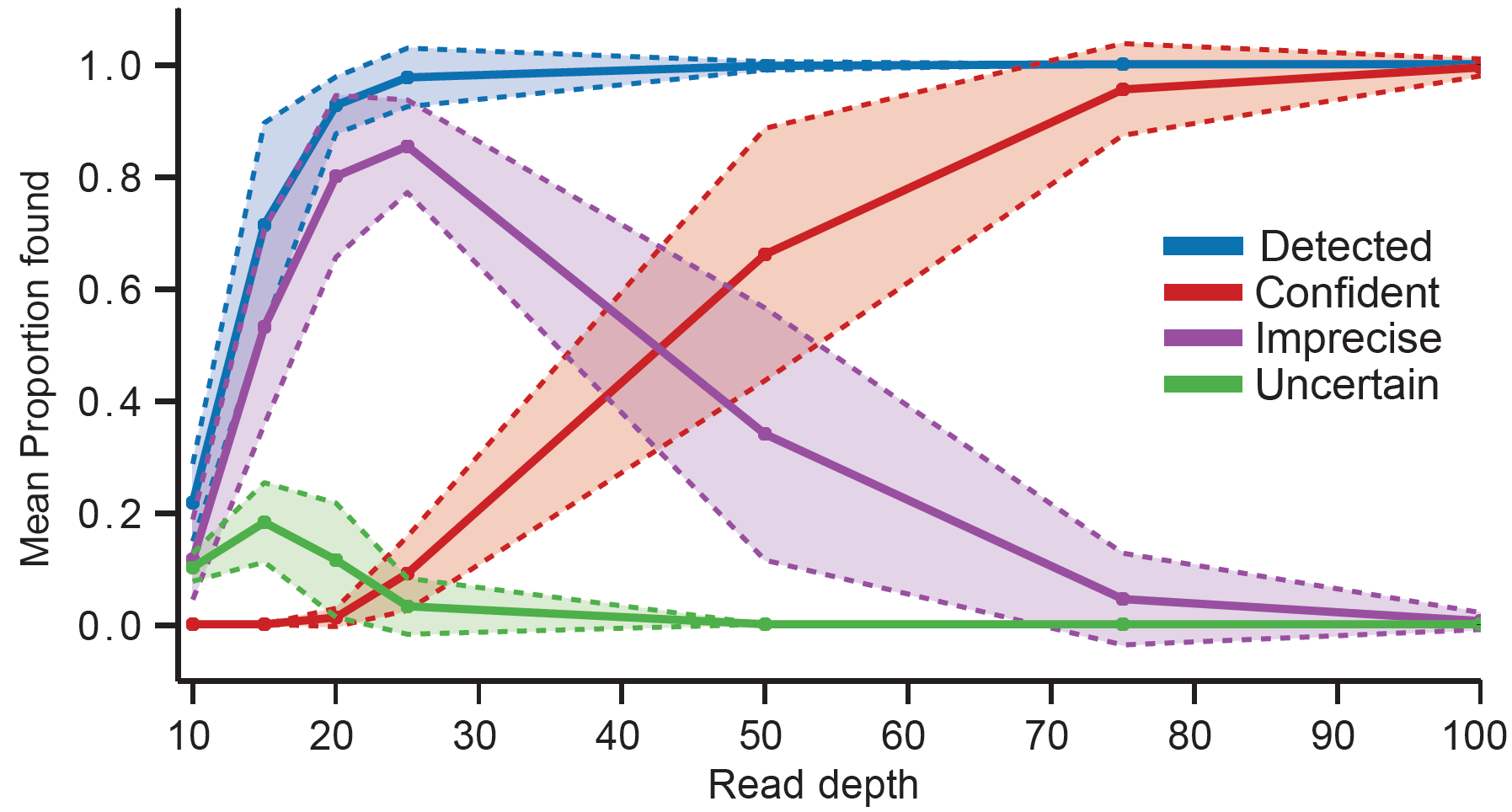
ISAba1 detection rate as a function of read depth, for seven *A*. *baumannii* GC1 genomes sequenced with Illumina. Mean (lines) and standard deviation (shaded areas) proportions of IS insertions correctly detected per genome, amongst 70 replicate read sets sampled at each depth level (7 read sets x 10 replicates each at read depths 10x, 15x, 20x, 25x, 50x, 75x, 100x). Blue, IS insertion site detected; red, detected with high confidence; purple, detected with low precision (larger than expected gap size between left and right flanking regions, indicated with ‘*’ in output); green, detected with low confidence (high-depth evidence for one side only, low-depth evidence for the other, indicated with ‘?’ in output).

### Comparison of ISMapper with TIF and *breseq*

The seven *A. baumannii* GC1 genomes were used to test both *breseq* [25] and TIF (Transposon Insertion Finder) [24]. *breseq* uses split read mapping to a reference genome along with statistical models to determine new junctions and deletions in the isolates of interest. As input, *breseq* takes paired end reads in FASTQ format, and a reference genome in Genbank format. The *breseq* manual indicates that new insertions of mobile elements can determined by looking for ‘JC JC’ evidence types in the final html output. All seven *A. baumannii* isolates were screened using default parameters and the reference genome A1 (accession CP010781). In all cases, *breseq* was unable to identify any mobile element insertions, including no structural variation at the known ISAba1 insertion sites, although many other types of structural variation were detected.

TIF requires as input paired end reads (FASTQ format), the head and tail sequences (approximately 17 bp) of the IS of interest as well as the size of the target site duplication the IS makes during transposition. TIF uses regular expressions to search for the head and tail sequences in the reads, and these reads are then extracted and grouped by their target site duplications. Unfortunately, following communication with the authors, we were unable to get TIF to output any results using our data. Other disadvantages of TIF are the requirements to (i) specify the size of target site duplications (which not all IS make and is not always known), (ii) manually extract subsequences of the IS rather than inputting the complete sequence, and (iii) manually edit a Perl script in order to specify inputs to the program.

### Example use case: Exploration of IS6110 insertions in *Mycobacterium tuberculosis*

While IS insertions are thought to be important for shaping the evolution of bacteria in a variety of ways, high-resolution comparative genomic studies of bacterial pathogens have largely ignored IS due to the difficulties associated with accurate detection of insertion sites from high-throughput short read data. An important example is IS*6110* in *M. tuberculosis* [34]. Profiling of IS*6110* insertions using PCR and restriction fragment based polymorphism (RFLP) based methods has been reported for typing purposes [35], and specific insertions have been linked to clinically relevant changes in function including in outbreak strains [9, 36, 37] However while numerous studies have reported the genomic analysis of hundreds of *M. tuberculosis* isolates sequenced on the Illumina platform, these have not included analysis of IS*6110* insertions. Thus, to demonstrate the utility of ISMapper for comparative profiling of IS in an important bacterial pathogen, we analysed the distribution of IS*6110* within 138 publicly available genomes representing the major lineages of *M. tuberculosis* [38]. Paired-end Illumina reads were downloaded from NCBI (ERP001731). A core genome phylogeny was generated from these reads by SNP (single nucleotide polymorphism) calling against reference genome H37Rv (accession NC_000962) (methods as described in [39]), followed by maximum likelihood phylogenetic inference on the SNP alignment using RAxML (GTR+G substitution model, 1000 bootstraps) to build a genome-wide phylogenetic tree. ISMapper was run with default settings to screen for insertions of IS*6110* (accession X17348) in each read set, relative to reference genome H37Rv.

A total of 392 unique IS*6110* insertion sites were identified by ISMapper, approximately one per 10 kbp of the 4.4 Mbp reference genome. The frequency of each insertion within each of the six main global lineages is shown in Figure 5b. The data indicate multiple lineage-specific IS*6110* insertions in lineages 2-6, but none that were shared by multiple lineages, suggesting that IS*6110* insertions began to accumulate only after *M. tuberculosis* diverged into these distinct lineages. Isolates in the “modern” lineages 2-4 and in the West African lineage 5 had more IS*6110* insertions overall, with far fewer insertions observed in the “ancient” East and West African lineages 1 and 6 (Figure 5c). Lineage 2, which includes the highly successful Beijing sublineage, had the highest number of IS*6110* although it was not the most common lineage in the collection (n=23); it could be that these insertions contribute to the adaptive fitness of the Beijing lineage.

**Figure 5:**
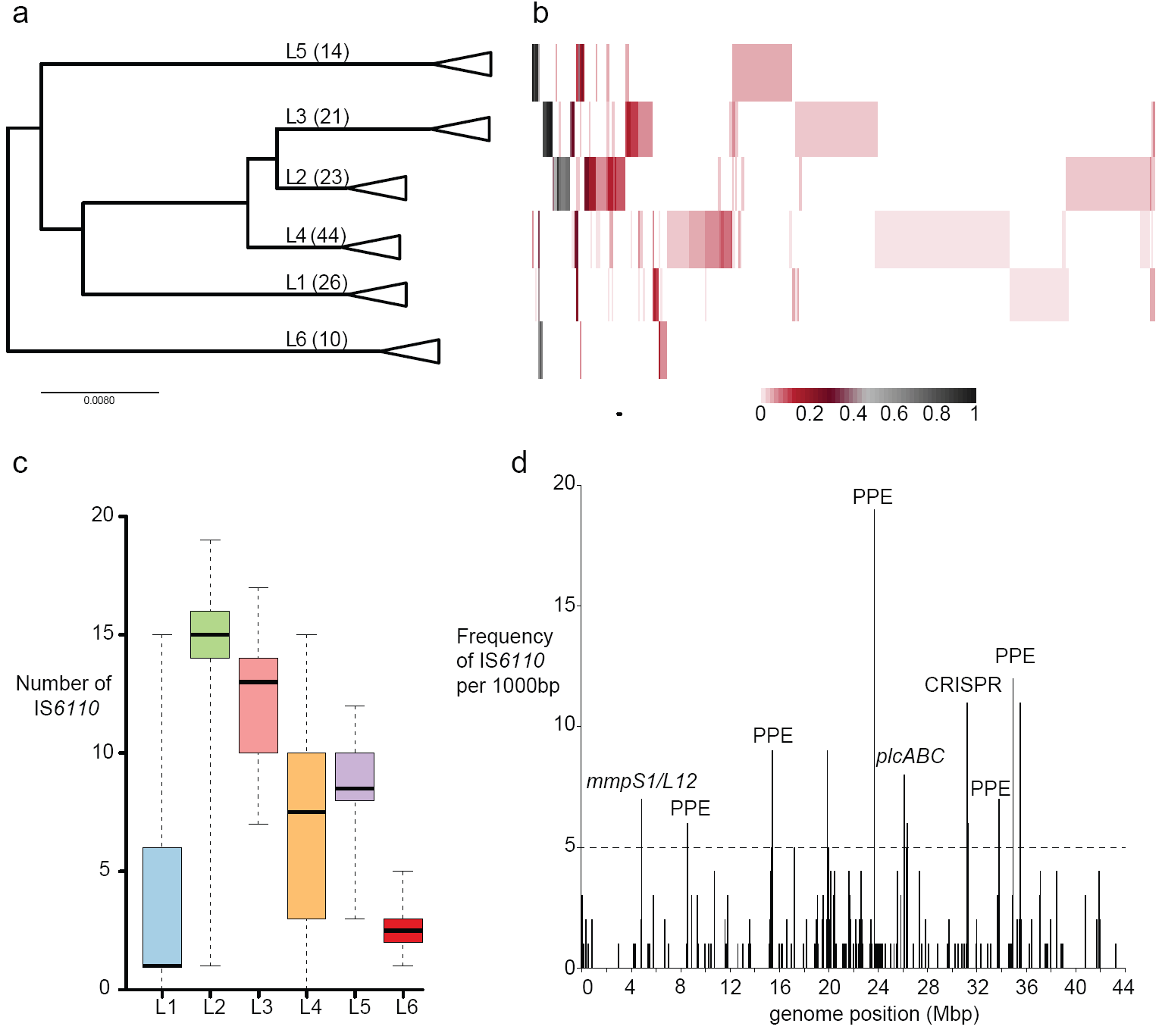
Analysis of IS*6110* insertions in a diverse set of *M. tuberculosis* genomes. Analyses are based on ISMapper analysis of publicly available Illumina paired end data from 138 genomes. (**a**) Phylogenetic tree for *M. tuberculosis* based on genome-wide SNP calls, with midpoint rooting and collapsed to lineage level. Number of isolates analysed in each lineage (L) is indicated in brackets. (**b**) Heat map lineage (rows), according to inset legend (i.e. a black cell indicates that the IS indicating the frequency of each IS*6110* insertion site (columns) detected in each insertion site was detected in all isolates of the given lineage, white cell indicates that the insertion site was not detected at all in that lineage). (**c**) Boxplots show number of IS*6110* insertions detected per genome, for each lineage. Black line, median; boxes, interquartile range; whiskers, minimum and maximum values. (**d**) Histogram of IS*6110* insertions in 1,000 bp windows along the *M. tuberculosis* chromosome. Dashed line, threshold for defining insertion hotspots.

The spatial distribution of unique IS*6110* insertions within the *M. tuberculosis* genome (Figure 5d) revealed several clusters of insertions detected by ISMapper. Many of these clusters comprised multiple independent insertions into PE/PPE genes (which are surface-associated and interact with the host immune system), as well as the membrane associated proteins *mmpS1* and *mmpL12*. There was substantial clustering of IS*6110* insertions interrupting genes encoding the CRISPR machinery, which is involved in immunity to bacteriophage and other foreign DNA. Further, all three phospholipase genes, which are involved in virulence by inducing cell death in macrophages [40] and are encoded by the *plcABC* operon, contained multiple IS*6110* insertions detected by ISMapper. This locus is a known hotspot for IS*6110* insertions and has been shown to mediate deletions of segments of this region [41]. IS*6110* insertions upstream of *phoP*, which have been associated with upregulation and enhanced virulence in *M. tuberculosis* [9], were identified in multiple lineages (1 insertion in 6 lineage 2 genomes; singular insertions in one genome each in lineage 3 and 5) and may be indicative of positive selection for enhanced *phoP* expression and virulence. These findings from ISMapper analysis are consistent with those reported from PCR-based screens of smaller sets of isolates, but provide a more comprehensive picture of IS dynamics in *M. tuberculosis* that could be extended to much larger genomic data sets and other important pathogens.

## Conclusions

ISMapper is a lightweight and reliable tool for the detection of IS insertion sites in bacterial genomes using high-throughput short-read sequencing data, which is now ubiquitous in microbial research and clinical investigations. ISMapper performed well on real and simulated data from 32 different IS and 13 bacterial species, detecting all but the most complex instances involving multiple neighbouring IS insertions or other repeated sequences. ISMapper was able to detect antimicrobial resistance-associated ISAba1 insertions in *A. baumannii*, with all sites detected by the program being subsequently confirmed by PCR. Compared to other tools such as *breseq* and TIF, ISMapper is ideal for detecting new positions for known IS in bacterial genomes. In addition, ISMapper was able to rapidly produce a wealth of data on IS*6110* insertions in *M. tuberculosis*, allowing quick identification of lineage-specific insertions and specific regions enriched for insertions that may be functionally significant.

## Availability and Requirements

- **Project name:** ISMapper
- **Project home page:** https://github.com/jhawkey/IS_mapper
- **Programming language:** Python v2.7.5
- **Operating system(s):** platform independent, requires Python 2.7 and dependencies
- **Other requirements:** BioPython v1.63, BWA v0.7.12, SAMtools v1.1, Bedtools v2.20.1, BLAST+ v2.2.28, Samblaster v0.1.21
- **License:** Modified BSD

## Authors’ Contributions

JH developed the code, analysed data and wrote the paper. KEH conceived the study and helped to draft the manuscript. HBJ participated in design and coordination of the study and contributed to data interpretation. RRW and DJE developed code. MH performed PCR and sequence analysis. RMH provided sequence data and isolates for validation and contributed to data interpretation. All authors read and approved the final manuscript.

## Acknowledgements

Australia (Fellowship #1061409 to KEH; Project Grant #1043830 to KEH and RMH) This work was supported by the National Health and Medical Research Council of and the Victorian Life Sciences Computation Initiative (VLSCI, #VR0082).

